# The development of navigation and attentional abilities shapes how children explore naturalistic environments

**DOI:** 10.64898/2026.09.29.755223

**Authors:** Qianqian Wan, Nathalie Moriarty, Shea E. Duarte, Simona Ghetti, Joy J. Geng

## Abstract

Exploration is fundamental to how children learn about the world. To explore a new environment comprehensively, a child must navigate without getting lost while simultaneously attending to salient information along the way. Both abilities improve across childhood, yet most studies examine them in isolation, often in constrained spaces with limited sensory complexity. Here, we asked 7- to 14-year-old children (N=42) and young adults (18-22; N=42) to freely explore a large furniture store in immersive virtual reality, which contained many objects and attention-capturing multisensory events. Surprisingly, despite traveling comparable distances, children explored less - they covered less spatial area and examined fewer unique objects than adults. Fine-grained analyses of their behaviors showed that the reduction in exploration was due to children spending more time backtracking and children taking longer to adapt to salient multisensory objects, which were uncorrelated across individuals after controlling for age and independently predicted exploration. However, only backtracking duration predicted success in a subsequent search task, demonstrating a link between navigational success during exploration and the quality of learned spatial representations for later use. Together, the results demonstrate that children explore less comprehensively than adults in large-scale spaces because they are susceptible to getting lost and perseverating on salient events nested within a large environment, but that these behaviors change with age. This suggests that while children are widely considered to be curious explorers within local environments, their ability to do so within larger environments is shaped by the maturation of cognitive systems that integrate experiences over complex contexts.

## 1. Introduction

Consider a child visiting an amusement park for the first time. To learn what the park offers, they must explore the park efficiently without getting lost while simultaneously processing unexpected, salient sensory events encountered along the way (e.g., a scream from a monster in a haunted house, music from a roller coaster, or lights from an ice-cream cart). For adults, navigating and attending to new information seems to occur without apparent costs, but for children, whose underlying systems are developing, these concurrent cognitive demands may prove challenging. Previous studies have characterized the development of children’s exploration of spatial environments by considering their navigation skills and describing how much ground they cover (Baumann et al., 2020; Farran et al., 2022; Heller et al., 2020; Saragosa-Harris et al., 2022; Šimon et al., 2019), or by the number of places and activities they engaged with (Babik et al., 2022; Gottlieb et al., 2020; Henderson et al., 1982; Ruff, 1984). However, each of these measures captures just one aspect of their exploratory experience, and most have been studied in relatively small spaces with few competing distractions. Yet navigation and attention to sensory information have almost never been studied together, and almost no work has asked what happens when the two must be coordinated within a single behavioral experience. We introduced children and young adults to a naturalistic, large-scale virtual environment containing many competing paths and sources of sensory information to ask a specific question: when navigation and attentional control are both developing and both demanded at once, do they represent a single dimension of developmental maturity, or do they leave independent behavioral signatures on how children explore?

From the earliest months of life, infants actively explore their surroundings, reaching for novel objects (Needham et al., 2002; von Hofsten, 2004; Yu & Smith, 2012), and directing their gaze toward novelty and complexity (Colombo & Mitchell, 2009; Dautriche et al., 2022; Kidd et al., 2012; Kwon et al., 2016). This early preference for novelty reflects a broader developmental pattern in which children tend to favor exploration over exploitation, sampling new information rather than processing familiar sources (Blanco & Sloutsky, 2024; Giron et al., 2023; Gopnik, 2020; Liquin & Gopnik, 2022). As children become mobile, this exploratory bias extends beyond gaze to encompass locomotion, with children traveling farther distances to investigate new locations and sample new information (Bramley & Ruggeri, 2022; De Simone & Ruggeri, 2022; Farran et al., 2022; Hills et al., 2015; Mehlhorn et al., 2015; Thurman & Corbetta, 2017). This coordination of sensory sampling and locomotion continues to develop through childhood and into adulthood (Lingwood et al., 2015, 2018; Newcombe, 2019, 2024; Newcombe & Shipley, 2015; Weisberg & Newcombe, 2016). Together, this work has led to the view that children are curious explorers who use their physical abilities to seek new information. However, these developmental patterns have been documented primarily in controlled settings with limited numbers of objects and relatively simple and small spaces. Whether children’s exploration remains as comprehensive in large-scale, navigable environments containing many competing multisensory events remains untested. This sets up two competing possibilities. If children are the curious, capable explorers that controlled studies suggest, they should explore these larger environments as comprehensively as adults. If comprehensive exploration instead depends on navigation and attentional abilities that are still developing, children should explore less comprehensively once a large space and competing sensory events demand both at once, even while remaining just as active. While exploration is often portrayed as effortless in laboratory tasks, this may not be true in naturalistic settings, which may depend on developing navigation mechanisms that support spatial memory and attentional mechanisms that regulate responses to salient information (Burles et al., 2020; Nazareth et al., 2018; Newcombe, 2019; Volkmer et al., 2022; Wetzel et al., 2021). Moreover, given how navigation and attention mechanisms have almost exclusively been studied in settings where only one is demanded at a time, it is unclear if developmental trajectories only become visible when both are simultaneously demanded. The typical controlled settings in laboratory task may systematically misrepresent how children explore the environments they actually inhabit.

Despite the natural combination of navigational and attentional demands during real-world experiences, most studies have focused on one or the other in isolation. Studies of map-based navigation show that efficiency improves over development (Burles et al., 2020; Nazareth et al., 2018; Newcombe, 2019). Older children with stronger navigation skills are less likely to become disoriented by spatial complexity, need fewer repetitions to learn, and make fewer errors in path reproduction compared to younger children (Brucato et al., 2022; Lingwood et al., 2015, 2018; Nys et al., 2015). Development of these skills allows children to explore more broadly and efficiently, covering more ground and building more accurate spatial memories of objects and places that can be used later (Brunec et al., 2023; Burles et al., 2020; Farran et al., 2022; Maier et al., 2025; Meade et al., 2019; Qin et al., 2014). In our hypothetical amusement park, older children would be expected to visit more rides throughout the park, build more robust cognitive maps while navigating, and retrieve a forgotten bag from a specific ride at the end of the day more efficiently. Notably, most tasks to date have used relatively low spatial demands, and the popular characterization of children as “natural explorers” may partly reflect the fact that spatial scales in laboratory and school-based tasks rarely challenge their developing spatial memory. Beyond the question of scale, developmental differences have been characterized almost exclusively through outcomes obtained after navigation (e.g., map accuracy, target retrieval), rather than during navigation itself. Whether children’s navigational ability has a measurable trace in the moment-by-moment record of exploration, and whether that trace predicts subsequent learning, remains largely unexamined.

Studies that have focused more on the development of attentional control show that older compared to younger children are more likely to reduce reflexive orientation towards perceptually salient stimuli as they learn to focus on more meaningful content (Blanco & Sloutsky, 2024; Corbetta & Shulman, 2002; DeBolt et al., 2023; Iarocci et al., 2009; Oakes, 2023; Posner & Rothbart, 2007; Ruff & Rothbart, 1996; Wan & Sloutsky, 2024). While surprising and salient events capture attention throughout the lifespan, the ability to learn from and adapt to them in order to reallocate attention to discover other objects and events improves with age (Blanco & Sloutsky, 2019, 2020; Fisher et al., 2014; Gopnik, 2020; Huang-Pollock et al., 2002; Wan & Sloutsky, 2024). Learning and adaptation ultimately result in more effective information sampling (Gottlieb & Oudeyer, 2018), comprehensive learning, and the resumption of ongoing goal-oriented behaviors (Sokolov, 1963; Wessel & Aron, 2017). If these developmental differences in attentional control extend to naturalistic exploration, children who adapt more slowly to repeated salient events should experience more interruptions to ongoing exploration, potentially limiting what they learn in the environment. Contrary to local notions of exploration, a strong attentional bias toward salient and novel events in children compared to adults may actually lead to less comprehensive global exploration if they fail to adapt to recurring salient stimuli. Together, the navigation and attentional control literatures have so far tested the effect of children’s developing cognition on behavior in constrained settings, precluding the ability to determine how they contribute to complex behaviors when both are simultaneously demanded.

We asked participants to freely explore a large, multiroom virtual furniture store containing ordinary objects and 10 multisensory objects that produced sounds and animations when approached (Figure 1). Here, we test the prediction that developmental differences in navigation and attentional control jointly shape exploration, such that children will cover less space and sample fewer objects than adults despite comparable movement. In a large, multiroom environment, navigation difficulty should be observable as prolonged retracing of familiar paths when spatial representations fail (Figure 2a), and developing attentional control should be observable as a reduced ability to adapt to recurring sensory events (Figure 2b). We also predict that these developmental signatures will affect the ability to remember the environment and locate objects in a subsequent task. Because both navigation ability and attentional control improve across this age range, a further question is whether they covary across individuals or represent separable developmental patterns. We focused on children ages 7 to 14 and young adults because gross motor systems are sufficiently mature to support large-scale exploration by early childhood (Adolph & Robinson, 2015), yet the cognitive systems underlying spatial memory and attentional control continue to improve into adolescence (Casey et al., 2005; Ghetti & Bunge, 2012; Keresztes et al., 2017).

**Figure 1.**
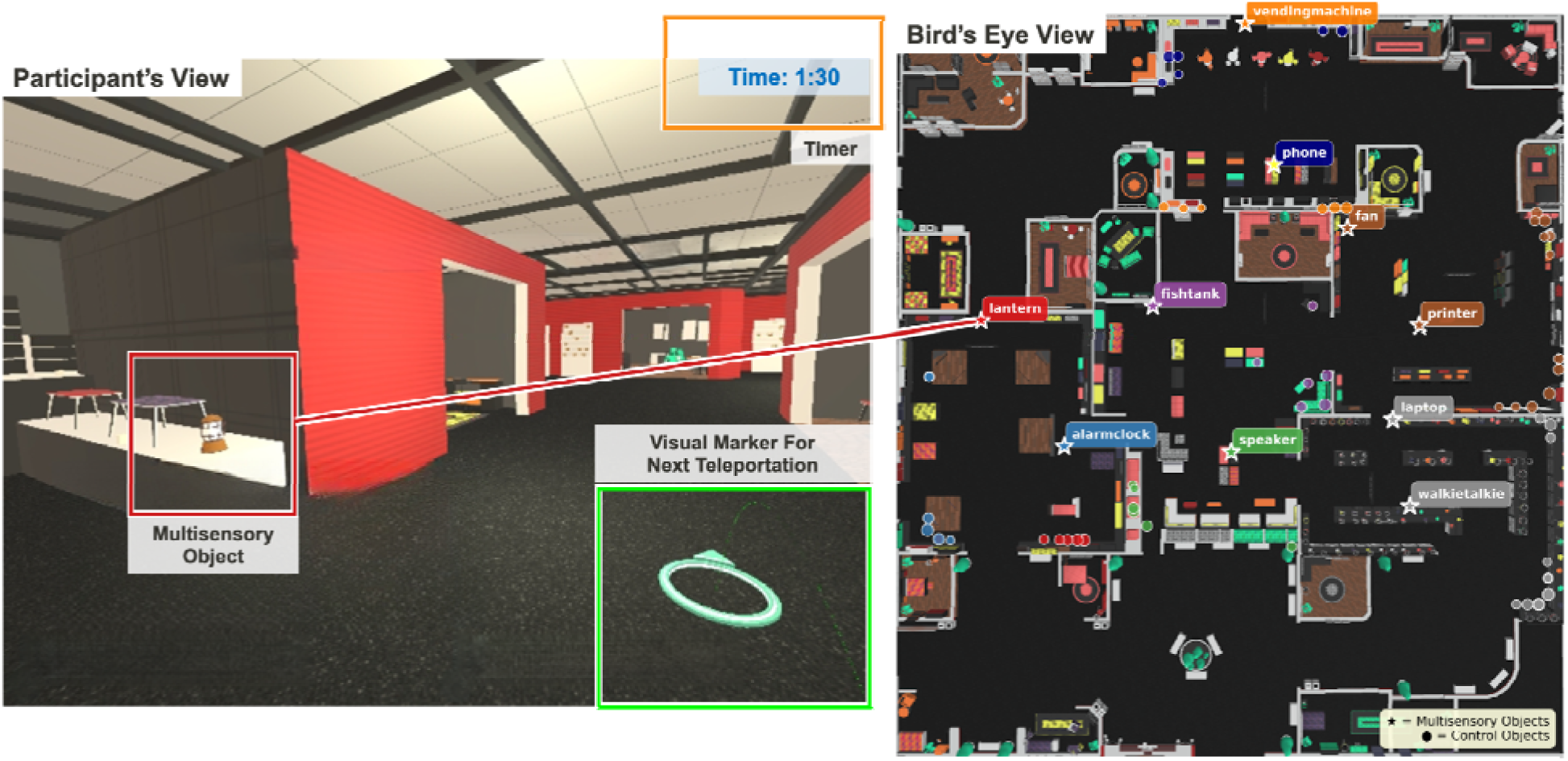
The virtual furniture store from the participant’s perspective and from a bird’s-eye view. The highlighted boxes in the left panel were not visible to participants but are used to highlight task features of a multisensory object (lantern in the red box), the teleportation marker (green box), and the elapsed time out of 4 minutes during the exploration period (orange box).

**Figure 2.**
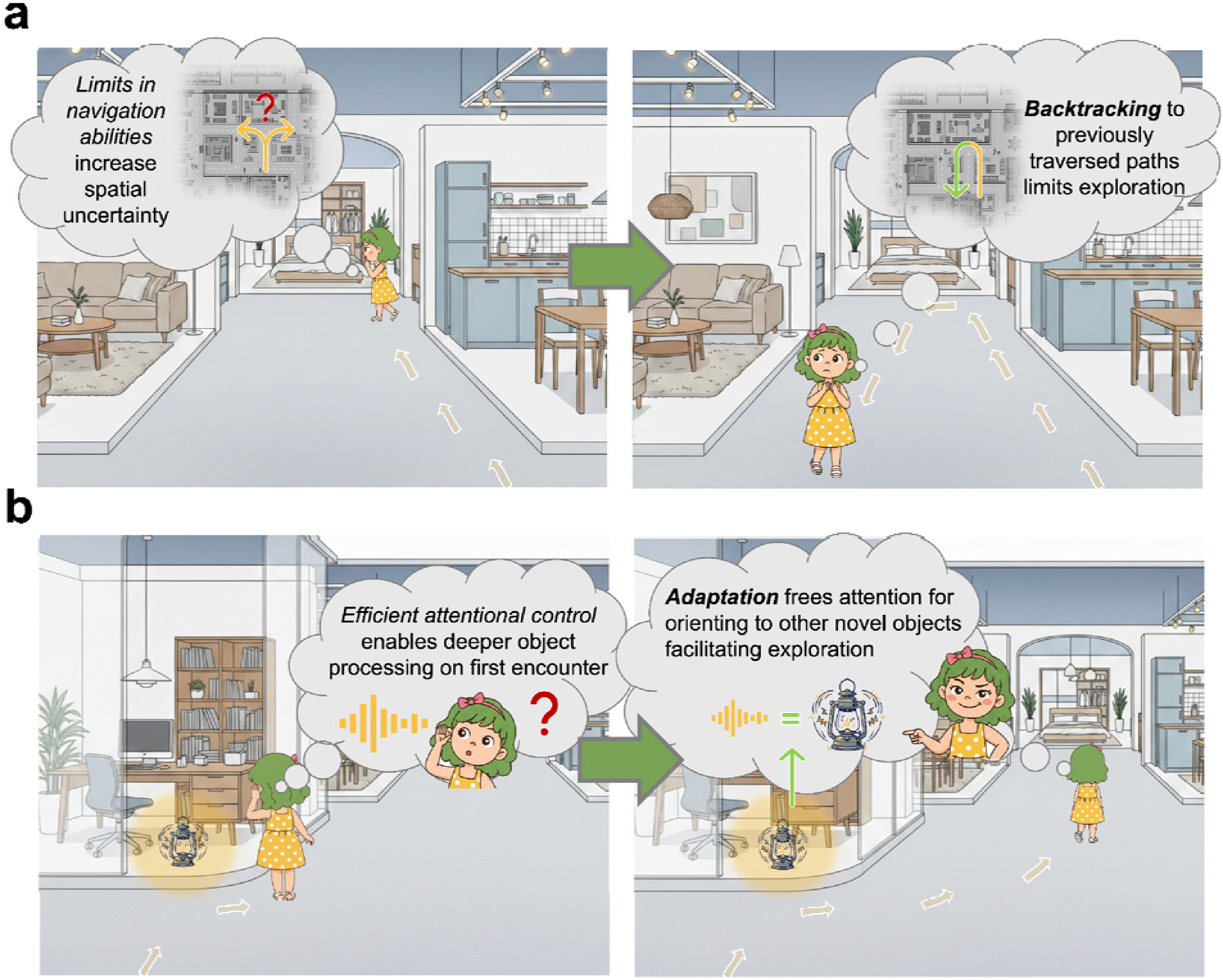
*Schematic illustration of the two key behavioral measures. (a) Backtracking duration is an index of navigation ability. During exploration, participants may experience uncertainty about their spatial position (left), leading them to backtrack for longer*, that is, to retrace previously traversed paths (right). The amount of time spent backtracking is taken as an index of the severity of navigational confusion. (b) Adaptation to multisensory events is an index of attentional control. When participants first encounter a multisensory object in a location (left), they orient attention toward the source and stop exploration. With repeated exposure to the same event (right), they are less likely to stop for it, reflecting learning about the object.

## 2. Method

### Participants

A total of 84 participants completed the study. Children (*N* = 42) ranged in age from 7.11 to 14.96 years (*M* = 11.10, SD = 2.11), and adults (*N* = 42) ranged from 18 to 23 years (*M* = 19.8, SD = 1.6). Of the children, 18 self-reported as girls and 24 as boys, and self-reported race and ethnicity included 20 White, 6 Multi-racial, 4 Latino, and 3 Asian participants. Of the adults, 24 self-reported as women and 18 as men; self-reported race and ethnicity included 23 Asian, 9 White, 3 Latino, and 1 Black participants. A sensitivity power analysis was conducted to determine the minimum detectable effect size given the sample sizes, alpha level, and desired power, independent of observed results ^71^. For age group comparisons (children: *n* = 41; adults: *n* = 42), two-sided independent-samples *t*-tests with α = .05 had 80% power to detect effect sizes of Cohen’s *d* = 0.62. Power calculations were performed using the statsmodels package in Python^72^.

Adult participants were recruited from the University of California, Davis, and received partial course credit in exchange for their participation. All adult participants reported normal or corrected-to-normal vision via contact lenses. Child participants were recruited from the developmental research subject pool and received a $15 gift card for their participation. All parents of child participants reported normal or corrected-to-normal vision.

To minimize the risk of cybersickness associated with head-mounted VR displays, all participants were screened for susceptibility to motion sickness prior to the study using the Visually Induced Motion Sickness Susceptibility Questionnaire^73^. Four adult participants who reported medium-to-low engagement and five children who did not complete the exploration phases were excluded, resulting in the final sample described above. All adult participants and parents of child participants provided informed consent, and child assent was obtained for all child participants, in accordance with the guidelines set by the University of California, Davis Institutional Review Board.

### Apparatus

Stimuli were presented on an HP Omnicept Reverb G2 head-mounted display (HMD) equipped with Tobii eye tracking. The HMD had a resolution of 4,320 × 2,160 pixels and a field of view of 114°. The eye tracker had a sampling rate of 120 Hz.

Participants completed the study in a sound-controlled data collection room. The HMD cable was tethered to a rod placed in the corner of the room to allow free body movement while participants stood at the center of the room. Participants navigated the virtual environment using teleportation and made behavioral responses using the HP Omnicept controller. The teleportation step was limited to 2 m to approximate a natural step size and prevent participants from teleporting long distances. The experiment was programmed using Vizard (WorldViz, Santa Barbara, CA) and run on SteamVR (Version 1.6.10, Valve Corporation, 2019) and Windows Mixed Reality.

### Virtual Environment

The virtual environment consisted of a 4,216-square-meter IKEA-inspired furniture store created in Vizard. The environment contained 23 rooms: 5 large showrooms and 18 smaller display rooms. Showrooms were large spaces featuring primarily one furniture type (beds, desks, couches, shelves, or lamps). Display rooms featured a variety of furniture arranged to represent functional settings (e.g., bedrooms, offices, living rooms). The remaining areas consisted of hallways decorated with plants, benches, and shelves. Doorways to each room were open, and some rooms had multiple entry points. The total environment contained 355 interactive objects (i.e., excluding structural elements such as floors, ceilings, and walls) available for visual exploration.

Ten multisensory objects were distributed across the environment (see Figure 1). These objects were designed to activate when the participant approached within 3 meters, producing sounds lasting 3 to 5 seconds, accompanied by corresponding animations of the same duration (e.g., a lamp that flickered and buzzed; Figure 1).

### Design and Procedure

#### Preparation Phase

Upon arrival, participants received instructions about VR safety, including staying within a designated area marked with tape. They were told they could turn around, crouch, or reach, but should only use teleportation to traverse the furniture store (Figure 1). Participants were also instructed to notify the experimenter immediately if they experienced any symptoms of motion sickness.

The HMD was fitted and adjusted for each participant. After fitting, a nine-point eye tracking calibration was performed, and the lens distance was adjusted for optimal viewing. Participants then completed a navigation practice session to familiarize themselves with the teleportation controls. Navigation was accomplished by pressing the thumbstick forward to aim at the desired location (maximum distance 2 m) and releasing to move there. Participants practiced until they demonstrated proficiency.

#### Exploration Task

The exploration task consisted of two 4-minute blocks. Participants began each block at the same starting position in the southwest corner of the environment. They were instructed to explore the space freely and learn it as thoroughly as possible. A timer displayed in the top corner of the HMD showed the remaining time (Figure 1).

#### Search Task

After completing both exploration blocks, participants received instructions for the search task. There were 10 search trials with five item types: bed, couch, lamp, shelf, and desk. Each target object was a piece of furniture located in the showrooms. On each trial, the target was presented in a blocked-off entryway of the virtual store for 6 seconds. Participants then navigated the furniture store to locate and select that object. Selection was accomplished by aiming a virtual laser beam using the controller and pulling the trigger. Targets could be selected from up to 1 meter away; the laser beam appeared red when the participant was too far from the object. If participants selected an incorrect object, they were informed of the error and could continue searching, with a 4-minute time limit per trial.

### Metrics

#### Exploration Coverage

Overall exploration of space and objects was assessed using spatial coverage and object coverage. Spatial coverage represents navigational exploration and was calculated as the proportion of the total area visited. The environment was divided into 100 equal grid cells, each approximately 6.23 by 6.84 meters, and coverage was defined as the number of cells entered during exploration. Object coverage represents attentional exploration and reflects the proportion of all 355 available objects that were fixated for longer than 300 ms during exploration.

#### Backtracking

To measure backtracking as a behavior reflecting navigation ability, we measured backtracking distance and backtracking duration from each participant’s navigation path. As illustrated in Figure 4a, we employed an algorithm to detect backtracking by first identifying sharp turns with an angle θ of 60° or less. The algorithm then evaluated whether the return path overlapped with the preceding forward path within a 5 m tolerance. For a segment to qualify as backtracking, the number of teleportation stops, n, had to be greater than 1, and each stop could span up to 3 m. The total backtracking distance was calculated as the sum of the distances (in meters) across all stops and detected backtracking instances and sessions. The total backtracking duration was calculated as the sum of the time spent on backtracking segments across sessions.

#### Adaptation to Multisensory Events

To measure adaptation to multisensory events as a behavior reflecting attentional control, we measured the mean stop duration following the onset of a multisensory object as an index of bottom-up interruption to ongoing exploration. All stop duration analyses focused on the 5 seconds following object activation and were baselined to the 5 seconds preceding activation. Stop durations were also compared with those following encounters with control objects, which were normal objects located in the same room as the multisensory objects (see Figure 1).

#### Search Performance

Following the exploration phase, participants searched for 10 target objects. Search time and success rate were measured across the remaining 9 trials, excluding the first target, which served as a practice trial.

## 3. Results

### Developmental Differences in Overall Exploration of Space and Objects

First, we quantified developmental differences in exploration within this large-scale naturalistic environment, using two metrics. Spatial coverage, our metric of navigational exploration, was calculated as the proportion of visited grid cells among the total 100 equal-sized cells. Object coverage, our metric of attentional exploration, was calculated as the proportion of total unique objects viewed.

As shown in Figure 3, consistent with the hypothesis that adults and older children explore more comprehensively, adults visited a higher proportion of total grid cells than children, *t*(82) = 3.93, M_diff_ = 0.114, 95% CI [0.056, 0.171], *p* < .001, Cohen’s *d* = 0.86. They also sampled a higher proportion of unique objects (i.e., furniture and decorative objects, *N* = 355), *t*(82) = 6.75, M_diff_ = 0.128, 95% CI [0.090, 0.166], *p* < .001, Cohen’s *d* = 1.47. Among children, the effect of age on spatial coverage was significant, *β* = 0.031, 95% CI [0.007, 0.055], *t*(40) = 2.64, *p* = .012, as was the effect of age on object coverage, *β* = 0.018, 95% CI [0.005, 0.031], *t*(40) = 2.81, *p* = .008, suggesting that both forms of exploration increase between 7 and 14 years. This pattern suggests exploration of both space and objects differs across development, with adults exploring more comprehensively than children, and age-related increases are evident within the child sample. However, the critical question we address next is what underlies these differences.

**Figure 3.**
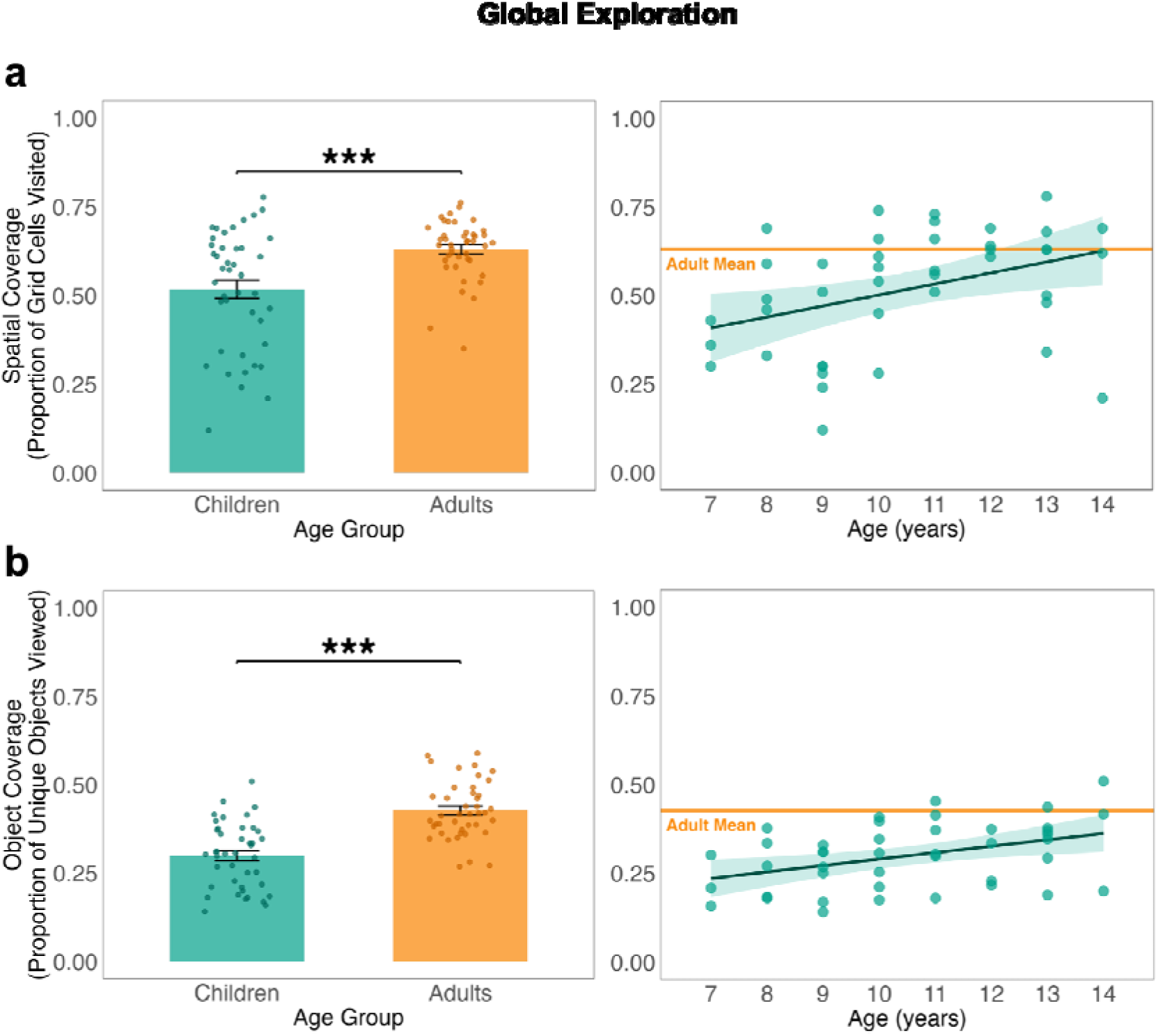
Two metrics of global exploration of the store by children and adults. (a) Proportion of total grid cells visited by children and adults (left) and age-related differences among children (right). (b) Proportion of unique objects viewed by children and adults (left) and age-related differences among children (right). The orange line represents the adult mean and serves as a reference against the range of performance in children across ages. Error bars represent standard error of the means.

### Backtracking as an index of navigation ability

One might expect that adults explore more simply because they have better motor skills that enable them to move faster. However, there were no differences in the absolute distance traveled between children (*M* = 369.72 m, SD = 132.93) and adults (*M* = 404.09m, SD = 128.46), *t*(81) = −1.20, M_diff_ = 38.99m, 95% CI [-31.13, 109.11], *p* = .235, Cohen’s *d* = −0.26. This rules out developmental differences in motor abilities for coordinating teleportation in VR or in the general speed of decision-making for movement.

During exploration, navigation difficulties can lead to feelings of being lost or uncertainty about one’s location. When this happens, navigators may try to regain their bearings by backtracking to an earlier position. We quantified backtracking in terms of the duration of a backtracking instance and its distance (See Figure 4a and Methods). We found that children (*M* = 86.41s, SD = 41.28) spent significantly more time backtracking than adults (*M* = 59.91s, SD = 41.20), *t*(82) = -2.94, M_diff_ = 26.50s, 95% CI [8.60, 44.40], *p* = .004, Cohen’s *d* = 0.64, as shown in Figure 4b. Among children, age was negatively associated with total backtracking duration (*β* = -6.42, 95% CI [-12.43, -0.40], *t*(39) = -2.15, *p* = .037). However, children (*M* = 121.21m, SD = 64.33) backtracked for a comparable distance to adults (*M* = 109.23m, SD = 78.56), *t*(79) = -0.76, M_diff_ = 11.99m, 95% CI [-43.15, 19.18], *p* = .447, Cohen’s *d* = 0.17. Among children, age did not significantly predict backtracking distance (*β* = 0.66, 95% CI [-9.25, 10.56], *t*(39) = 0.13, *p* = .894). The difference in backtracking duration across age, but not in backtracking distance, suggests that children backtracked because they were confused about their current location and needed extra time to determine their location relative to the starting point.

Further support for this interpretation comes from the speed profile during backtracking segments. Approach speeds during forward exploration were statistically identical between children and adults (M_adults_ = 2.09 m/s, M_children_ = 2.16 m/s, 95% CI [−0.55, 0.33], Cohen’s d = −0.11), ruling out general differences in movement speed. However, return speeds during backtracking were significantly lower in children than adults (M_adults_ = 2.03 m/s, M_children_ = 1.73 m/s, 95% CI [0.03, 0.91], Cohen’s d = 0.45). Children also showed a greater asymmetry between their approach and return speeds than adults did.

### Adaptation to multisensory events as an index of attentional control

Children may also have explored less because of their responses to salient information in the environment. Several multisensory, dynamic objects were interspersed throughout the space, likely capturing attention. We quantified participants’ experiences with these objects as an indicator of attentional control during exploration.

Adaptation to multisensory events was measured by the duration of an individual’s pause immediately after the onset of a surprising sound across three encounters. Because the sound was triggered only when an individual traveled near the object, individuals encountered a different number of unique multisensory objects (*M* = 6 out of 10) and for a different number of times (*M* = 2.5). To study how participants might adapt to these multisensory objects, we included only the first 3 encounters with each unique object to ensure sufficient power. Behavior in response to the multisensory objects was compared with that to control objects located in the same rooms (see Methods).

As illustrated in Figure 5, a linear mixed-effects model predicting mean changes in stop duration revealed a significant three-way interaction between age group × event type (multisensory, control) × number of encounters (*β* = 0.447, 95% CI [0.079, 0.904], *t*(2475)^1^ = 3.18, *p* = .002). Adults showed a significant decrease in average stop duration to multisensory events across encounters (encounter 1 vs 2: *t*(2326) = 1.58, *p* = .256, 95% CI [-0.11, 0.58]; encounter 1 vs 3: *t*(2343) = 4.59, *p* < .0001, 95% CI [0.54, 1.65]; encounter 2 vs 3: *t*(2325) = 3.36, *p* = .002, 95% CI [0.26, 1.46]), but not to control events (*p*s > .179). The average stop duration to the multisensory event differed from the control object at encounter 1 (*t*(2317) = -8.13, *p* < .0001, 95% CI [-1.11, -0.68]) and encounter 2 (*t*(2310) = -2.15, *p* = .0314, 95% CI [-0.74, -0.03]), but not at encounter 3 (*t*(2321) = 1.01, *p* = .3128, 95% CI [-0.31, 0.95]), indicating that adult stop durations decreased over subsequent encounters with the third encounter being equivalent to control objects. In contrast, children did not show significant adaptation; there were no differences in mean stop duration across encounters (encounter 1 vs 2: *t*(2307) = 0.48, *p* = .881, 95% CI [-0.31, 0.47]; encounter 1 vs 3: *t*(2338) = 0.34, *p* = .937, 95% CI [-0.49, 0.66]; encounter 2 vs 3: *t*(2323) = 0.013, *p* =1.00, 95% CI [-0.61, 0.62]). There was no significant main effect of age (*β* = 0.261, 95% CI [-0.107, 0.630]), *t*(979) = 1.39, *p* = .164) suggesting that the main developmental difference was the lack of adaptation across encounters in children, rather than a difference in stop duration. Critically, this pattern of results was observed despite adults initially spending comparable, if not more, time processing multisensory events to children (*M*_adult at encounter 1_ - *M*_children at encounter 1_ = 0.25s, 95% CI [-0.01, 0.51], *t*(411) = 1.86, *p* = .063). Thus, the rapid adaptation across repetition in adults was not due to lower initial attentional orientation. A supplementary analysis showed that this pattern was magnified when participants fixated on the sound source (see Supplementary Materials).

**Figure 4.**
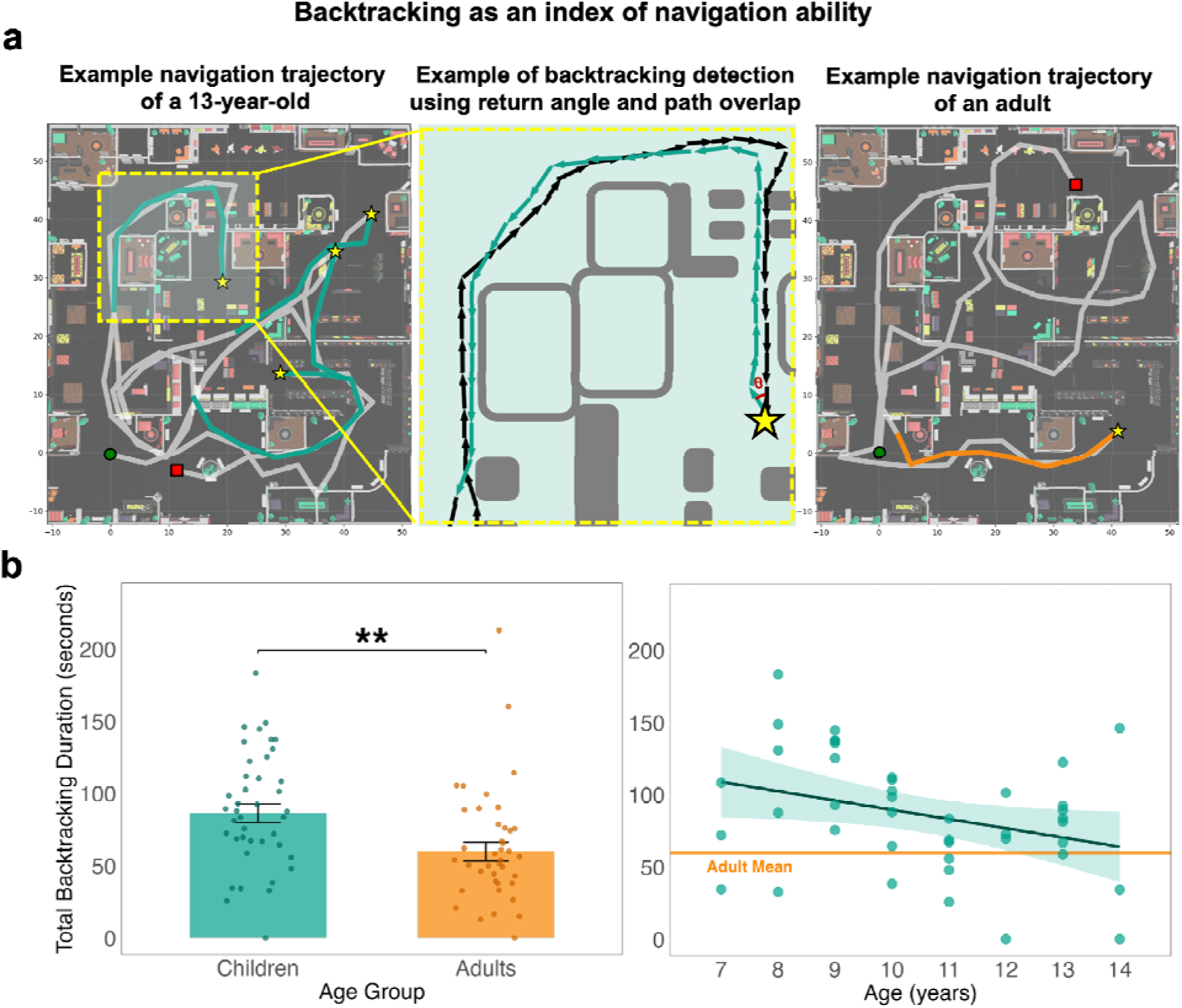
Backtracking detection method and age-related differences in backtracking duration. (a) Example navigation paths from a 13-year-old (left) and an adult (right), and a schematic illustration of backtracking path detection (middle). Notes. Yellow stars indicate turning points. The green circle marks the starting position and the red square marks the ending position. For a portion of the path to be classified as backtracking, the return angle θ must be equal to or smaller than 60°, and the number of teleportation stops (as shown in green arrows) must be greater than 1. The return path must overlap with the forward path within a tolerance of 5 m. Total backtracking distance is the sum of the distances of all stops backtracked across sessions. Total backtracking duration is the sum of the time a participant spent backtracking across sessions. (b) Total backtracking duration in seconds by children and adults (left) and age-related differences among children (right). Error bars represent standard error of the means.

**Figure 5.**
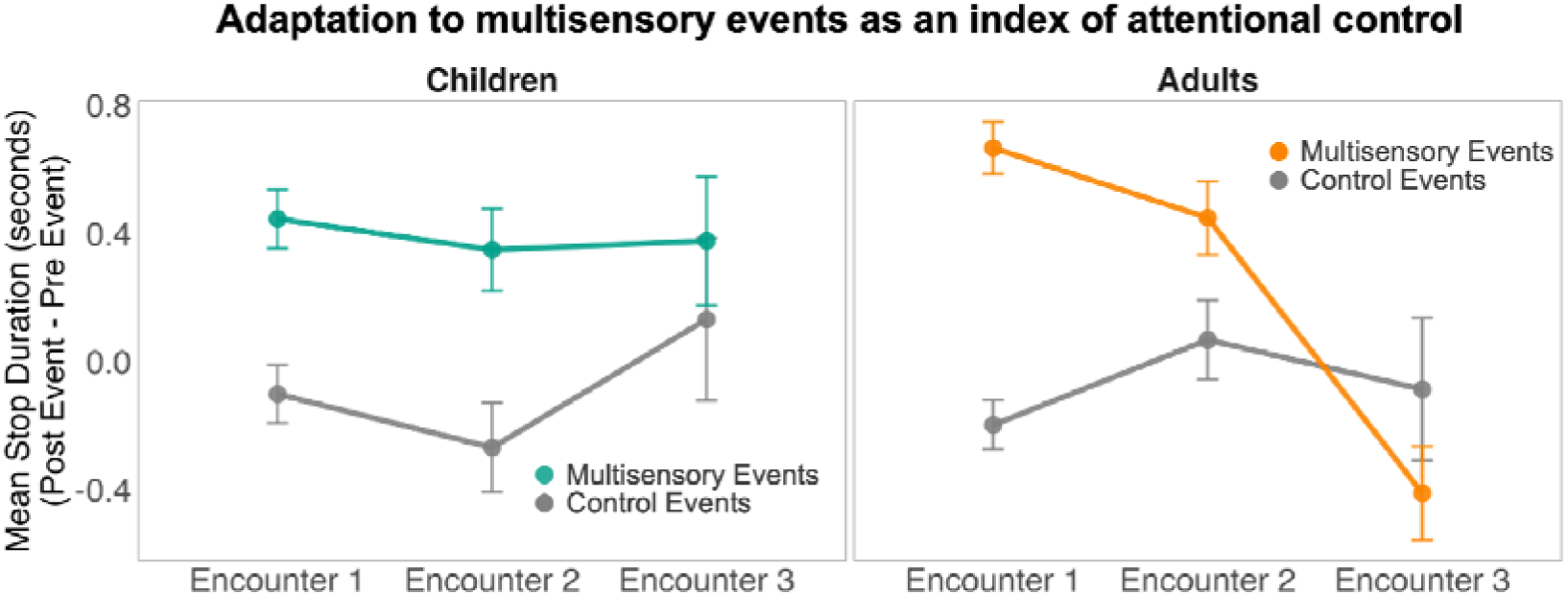
Average stop duration in response to multisensory and control events in children and adults during the first three encounters with the same object. The mean stop duration in response to the multisensory or control event was calculated by subtracting the stop duration from the 5-second period prior to the onset of the multisensory event (the pre-event) from the 5-second period after the event was triggered (the post-event). This normalization of the post-event data was necessary to account for general inter-individual variability in stop durations (see Methods). Error bars represent standard error of the means.

Within the child sample (ages 7 to 14; 1052 events), there was a significant main effect of multisensory events on average stop duration (*β* = 1.70, 95% CI [0.212, 3.180], *t*(1001) = 2.24, *p* = .025), but no interaction with age. Children stopped for longer when encountering multisensory events, consistent with attentional capture, but this did not change with age. Overall, adults and children initially paused for comparable durations in response to multisensory events, but adults rapidly adapted across encounters while children did not. The absence of adaptation in children suggests that attentional control over salient environmental information continues to develop through childhood.

### Backtracking and adaptation to multisensory events make independent contributions to exploration

Before looking at the joint effect of backtracking and multisensory adaptation on exploration, we first considered their contributions independently. When tested individually, backtracking duration predicted spatial coverage in children (*β* = -0.002, 95% CI [-0.00264, -0.00078]), *t*(80) = -3.64, *p* < .001, but not in adults (*p* = .943, 95% CI [-0.00097, 0.00090]), and had no significant effect on object coverage (*p*s > .09, largest *β =* -0.00056, 95% CI [-0.00119, 0.00008]). Adaptation predicted object coverage in children (*β* = -0.044, 95% CI [-0.0745, -0.0138], *t*(74) = -2.90, *p* = .005) but not in adults (*p* = .607, 95% CI [-0.028, 0.047]), and had no significant effect on spatial coverage (*p*s > .714, largest *β* = -0.015, 95% CI [-0.094, 0.064]). The correlation between backtracking and adaptation, accounting for age and number of multisensory events encountered, was not significant, r(76) = 0.026, 95% CI [-0.199, 0.248], p = 0.824. This lack of a correlation is itself a substantive finding: two behavioral measures that both improve with age and both predict exploration outcomes share no variance across individuals, indicating that navigation ability and attentional control represent separable dimensions of developing cognition rather than a single underlying maturational factor.

However, because navigation and attention operate concurrently during exploration, each measure’s full contribution may only be visible when the other is accounted for. We therefore used model comparison to determine the relative contributions of each to spatial and object coverage, our metrics of navigational and attentional exploration. We fit a series of regression models with age group, backtracking, and adaptation to multisensory events as predictors across all possible combinations (see Supplementary Tables S1 and S2 for full model lists and fitting results). The best-fitting model was selected based on AIC, BIC, and adjusted R², with priority given to the model favored by the majority (2 out of 3) of the criteria when they disagreed.

The best-fitting model for spatial coverage was the full model, which revealed significant interactions between backtracking duration and age group (*β* = -0.0019, 95% CI [-0.00325, -0.00056], *t*(71) = -2.83, *p* = .006) and between adaptation to multisensory events and age group (*β* = -0.19, 95% CI [-0.348, -0.027], *t*(71) = -2.33, *p* = .023). Both backtracking duration and adaptation predicted spatial coverage in children (β_backtracking_ = -0.0024, 95% CI [-0.00328, -0.00147], *t*(71) = -5.26, *p* < .001); *β*_adaptation_ = -0.16, 95% CI [-0.263, -0.066], *t*(71) = -3.34, *p* = .001, but not in adults (*β*_backtracking_ = -0.0005, 95% CI [-0.00147, 0.00052], *t*(71) = -0.944, *p* = .348; *β*_adaptation_ = 0.023, 95% CI [-0.104, 0.150], *t*(71) = -0.36, *p* = .723). Notably, adaptation emerged as a predictor of spatial coverage only in the joint model. This model explained 38.1% of the variance, *F*(7, 71) = 7.86, *p* < .001. The main effects of age group, backtracking duration, and adaptation were not significant, *p*s > 0.1.

For object coverage, the best fitting model included main effects of backtracking duration (*β* = -0.007, 95% CI [-0.00115, -0.00026], *t*(74) = -3.17, *p* = .002,), adaptation to multisensory events (*β* = 0.017, 95% CI [-0.020, 0.054], *t*(74) = 0.917, *p* = .362), and age group (*β* = -0.11, 95% CI [-0.148, -0.070], *t*(74) = -5.59, *p* < .001), as well as an interaction between age group and adaptation (*β* = -0.057, 95% CI [-0.105, -0.009], *t*(74) = -2.38, *p* = .020). The main effect of backtracking showed that individuals who spent more time backtracking, suggesting greater navigational uncertainty, viewed fewer unique objects. More interestingly, the interaction showed that adaptation predicted object coverage in children (*β* = -0.04, 95% CI [-0.0699, -0.0101], *t*(74) = -2.67, *p* = .009) but not adults (*β* = 0.017, 95% CI [-0.020, 0.054], *t*(74) = 0.917, *p* = .362). Thus, children who adapted more to multisensory events sampled more objects overall, suggesting that developing attentional control contributes to greater object coverage. The model explained 42.8% of the variance, *F*(4, 74) = 15.62, *p* < .001.

### Backtracking during exploration predicts subsequent spatial memory for object locations

Following the eight-minute exploration phase, participants searched for 10 target objects in the environment. The visual search trials provide a test of the practical value of behaviors during the exploratory period for subsequent behaviors in the same environment. Overall, as shown in Figure 6, adults (*M* = .848, SD = .136) found more targets than children (*M* = 0.745, SD = 0.208), *t*(81) = 2.67, M_diff_ = 0.102, 95% CI [0.026, 0.179], *p* = 0.009, Cohen’s *d* = 0.59. The effect of age within children, *β* = 0.060, 95% CI [0.034, 0.086], *t*(39) = 4.6, *p* < 0.001, suggests an improvement in search success between 7 and 14 years. However, total search times for successful trials were similar between children and adults (*M*_children_ = 37.99s, SD = 17.52s; *M*_adult_ = 33.00s, SD = 11.63s), *t*(81) = 1.53, M_diff_ = 4.99s, 95% CI [-1.49, 11.47], *p* = 0.129, Cohen’s *d* = .34. When children did locate a target, they navigated to it as quickly as adults, indicating that the bottleneck was not motor execution or general decision speed but the quality of the spatial representation built during exploration. This closes a direct chain from navigational confusion during exploration, indexed by backtracking duration, through reduced spatial coverage, to impoverished spatial memory for object locations.

**Figure 6.**
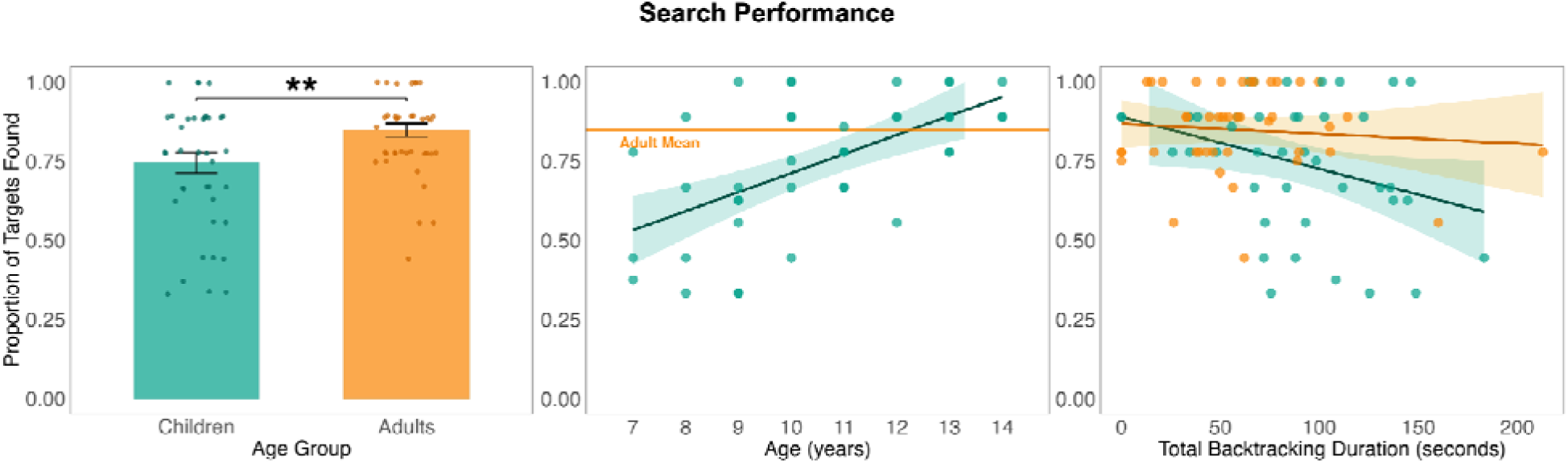
Search performance by children and adults (left), age-related differences among children (middle), and correlation with total backtracking duration (right). Error bars represent standard error of the means.

To determine whether backtracking duration and adaptation to multisensory events during exploration predicted subsequent search success, we fitted models of search success rate with backtracking duration, adaptation to multisensory events, age group, and all possible interaction terms as predictors (see Supplementary Table S3). Using the same model comparison criteria, the best-fitting model explained 23.8% of the variance and included an interaction between age group and backtracking duration (*β* = -0.002, 95% CI [-0.00392, -0.00047], *t*(74) = -2.17, *p* = .013). Children had a steeper slope (*β* = -0.0025, 95% CI [-0.00378, -0.00122], *t*(34) = -3.08, *p* = .001) than adults (*β* = -0.0003, 95% CI [-0.00146, 0.00085], *t*(40) = -0.59, *p* = .556), indicating that children who backtracked more were less likely to find targets, whereas this was not the case for adults (Figure 6, right). This pattern suggests that backtracking duration, reflecting navigation ability, predicted not only spatial coverage during exploration but also how effectively children could later locate objects in the environment. The main effects of age group and backtracking duration were not significant, *p*s > .392.

## 4. Discussion

Children aged 7 to 14 explored a large naturalistic environment just as actively as adults, covering comparable distances, yet learned less from that exploration. Two behavioral signatures, backtracking duration and adaptation to repeated multisensory events, explained this gap and, critically, did so independently of each other, pointing to separable rather than unified developmental patterns in exploratory learning.

### Backtracking duration and adaptation to multisensory events capture distinct developmental signatures during exploration

Backtracking duration captures navigational confusion as it unfolds during exploration, rather than inferring it from a downstream memory test. Children backtracked over similar distances as adults but spent significantly longer doing so, and this duration decreased with age. If children were simply choosing to retrace a familiar route as a navigation strategy, we would expect their backtracking distance and duration to covary. Instead, this developmental dissociation reveals that children paused and reoriented in real time because they were uncertain about their current location, a reading reinforced by the speed profile: approach speeds were identical across age groups while return speeds were selectively lower in children, producing an approach-to-return asymmetry consistent with uncertainty-driven hesitation rather than deliberate strategic retracing. This interpretation aligns with developmental changes in spatial representations, as children in this age range transition from egocentric, landmark-based strategies toward flexible allocentric representations that support cognitive maps (Burles et al., 2020; Ekstrom & Hill, 2023; Farzanfar et al., 2023; Nys et al., 2015), supported by the gradual maturation of hippocampal and retrosplenial systems (Newcombe, 2019; Wills et al., 2014). Backtracking duration predicted spatial coverage and, independently, object coverage in children but not adults, with the latter likely reflecting fewer opportunities to encounter unique objects when spatial coverage was limited. This process-level signature also carried forward: children who backtracked more found fewer targets in the subsequent search task, yet when they did locate a target they navigated to it as quickly as adults, indicating that the bottleneck was the quality of the spatial-categorical map built during exploration, specifically knowledge of which region of the environment housed which type of furniture.

Adaptation to multisensory events reflects the development of attentional control. Adults initially stopped for longer when encountering multisensory events than children did, but adults diminished their stops quickly with repeated exposure, whereas children showed sustained, moderate responses across all three encounters. Adaptation predicted object coverage in children but not in adults, consistent with the interpretation that developing attentional control shapes how broadly children sample objects during exploration. Although children as a group did not show adaptation, there were meaningful individual differences such that children who adapted faster also sampled a greater proportion of unique objects. More surprisingly, adaptation also predicted spatial coverage in children but not in adults when both behavioral measures were modeled jointly, suggesting that the ability to adapt to recurring sensory events extends to how broadly children explore space. One might suspect that these associations simply reflect time diverted from exploration. However, children did not spend overall more time stopped at multisensory objects than adults did. What differed was how stop durations changed across encounters with the same object, suggesting that the adaptation effects reflect a more general ability to learn from and remember sensory experiences within their contexts over time. Evidence that salient distractors can be rapidly ignored after only a few trials has been reported in adults (Won et al., 2022; Zhang & Gaspelin, 2026), but the current results show that adaptation can occur on a much longer timescale with potentially interfering events between encounters. The supplementary fixation analysis supports this interpretation and points to a specific locus for the developmental difference. When adults fixated the sound source, their initial stops were longer and their subsequent adaptation was steeper, suggesting that visual contact with the source deepened the initial encoding in a way that facilitated later suppression. Children who fixated showed longer initial stops but did not show complementary steeper adaptation across encounters, indicating that the developmental difference lies not in whether children oriented to the event but in whether that orientation produced an encoded representation robust enough to support subsequent suppression. This pattern extends the laboratory literature on attentional development (Blanco & Sloutsky, 2019, 2020; Fisher et al., 2014; Gopnik, 2020; Huang-Pollock et al., 2002; Volkmer et al., 2022; Wan & Sloutsky, 2024; Wetzel & Schröger, 2007; Wetzel et al., 2011, 2021) into a naturalistic context where the consequence of slower adaptation was not simply a longer reaction time on a single trial, but a reduction in the breadth of information sampled across the entire exploration period.

### Scale and concurrent cognitive demands reveal developmental differences that controlled tasks cannot capture

The introduction raised the possibility that comprehensive exploration requires the coordination of navigation and attention, that this coordination is itself developmental, and that both scale and concurrent sensory demands are necessary to reveal this. Our findings are consistent with both claims. The lack of a significant correlation between backtracking and adaptation, after accounting for age and number of multisensory events encountered, indicates that navigation ability and attentional control represent separable dimensions of development, each contributing independently to exploration outcomes, consistent with the view that these measures tap distinct processes that co-occur during naturalistic exploration.

The spatial scale of the environment was necessary to reveal a signature behavior of children’s navigational ability. In smaller environments where maintaining spatial orientation is relatively easy, navigation ability is unlikely to leave a measurable trace during exploration itself. In isolated navigation tasks, children’s difficulties typically manifest as discrete errors such as wrong turns, pauses, and less systematic coverage. The prolonged backtracking we observed is qualitatively different, reflecting sustained spatial confusion that unfolds continuously across an exploration session rather than momentary navigational errors at discrete choice points (Farran et al., 2022; Lingwood et al., 2015, 2018; Nys et al., 2015).

The concurrent presence of multisensory events within a larger navigational task was necessary to reveal the costs of attention to saliency. In laboratory paradigms where attention is the sole demand, children show strong initial orienting and can adapt to repeated novel events 35. In our environment, adaptation had to occur over longer delays in repeated exposure. Under these conditions, a clear developmental pattern emerged: adults, who routinely navigate large, sensory-rich environments in daily life, efficiently adapted their attentional responses and moved on, whereas younger children, for whom environments at this scale and sensory density may be less familiar, showed less labile adaptation. The most parsimonious account of why adaptation predicted spatial coverage only when backtracking was included in the statistical model is that backtracking durations were overall longer and more variable across individuals and when unaccounted for, created variance that obscured the comparatively smaller contribution of stop durations at multisensory events. This does not require assuming that navigation and attention interfere with each other at a cognitive level, since the two were not correlated, but only that both backtracking and adaptation contribute to the total exploration time. Together the results demonstrate that at the scale of everyday environments such as amusement parks, museums, or schools, where navigation and attention to sensory events co-occur, what children learn from exploration is shaped by both the spatial representations they are able to build and by how effectively they regulate their attentional processing.

### Limitations and Future Directions

The present study identified two behavioral measures that predict exploration outcomes and offered cognitive interpretations grounded in the existing literature. However, our account is necessarily incomplete. Exploration in naturalistic environments likely depends on additional cognitive processes beyond spatial memory and attentional adaptation. Executive functions such as planning, which develop substantially across this age range (Diamond, 2013), may influence how children select among available paths and prioritize certain areas of the environment. Curiosity and intrinsic motivation, which vary both across development and across individuals within an age group (Gruber et al., 2014; Kidd & Hayden, 2015), could shape how long children choose to remain in a given area and how willing they are to leave a known region for an unfamiliar one.

Additionally, the exploration task was open-ended with no explicit goal. Therefore, in its current form, it cannot be directly used to test whether the exploration-exploitation trade-off shifts in naturalistic environments, as laboratory studies suggest (Blanco & Sloutsky, 2024; Gopnik, 2020; Hilton & Sabbagh, 2025; Liquin & Gopnik, 2022). Our findings do establish that developing navigation ability and attentional control shape exploration itself, even when the task affords nothing but exploration. Future work could build computational models on the behavioral metrics developed here to formalize how these cognitive signatures interact with goal states to shift the balance between exploration and exploitation across environments and development.

The independence of these two signatures raises a question for future work of whether their separability holds across environments that vary in spatial scale and sensory density, and whether the time costs each imposes on exploration change as environments become more or less demanding along each dimension. Future work tracking the sequencing of navigation and attentional events within individual paths could also address whether moments of spatial confusion systematically precede or follow encounters with salient sensory events, providing a finer-grained test of how these two processes unfold in relation to each other during exploration.

Finally, teleportation-based VR navigation lacks the proprioceptive and vestibular feedback of physical walking. Whether the same behavioral signatures and their relations to exploration outcomes would be observed during fully embodied navigation in real environments remains an open question.

Children explore less comprehensively than adults not because they move less, but because two developing cognitive systems are each still maturing, each leaving its own trace on how children navigate the environment and regulate their responses to recurring events. This approach moves beyond documenting that exploration improves with age toward identifying the specific processes that drive that improvement, showing that their concurrent operation in rich, large-scale environments produces a developmental trajectory that neither process produces when studied in isolation.

## Acknowledgments

We would like to thank Rishit Das, Eliana Ertsy, Zoe Hareng, Makayla Souza-Wiggens, Catherine Halpern, and Morgan Peters for help with data collection and thoughtful discussions. This work was funded by an award from the James S. McDonnell Foundation to JG and SG.

## Data Availability Statement

All data and analyses reported can be found on OSF: https://osf.io/wn4b9/overview?view_only=06f53f6e4743489da0af12ed47b4f12c/.

## Supplementary Information

**Table S1.** Model Comparison for Spatial Coverage Analysis.

| Model | Param No. | AIC | BIC | Adj. R<sup>2</sup> | Formula |
| --- | --- | --- | --- | --- | --- |
| 2 | 2 | -98.00 | -93.26 | 0.188 | Spatial Coverage = Total Backtracking Duration |
| 3 | 2 | -81.25 | -76.51 | -0.003 | Spatial Coverage = Adaptation to Multisensory Events |
| 4 | 2 | -95.50 | -90.76 | 0.162 | Spatial Coverage = Age Group |
| 5 | 3 | -96.21 | -89.10 | 0.180 | Spatial Coverage = Total Backtracking Duration + Adaptation to Multisensory Events |
| 6 | 3 | -105.51 | -98.41 | 0.271 | Spatial Coverage = Total Backtracking Duration + Age Group |
| 7 | 3 | -93.50 | -86.40 | 0.151 | Spatial Coverage = Adaptation to Multisensory Events + Age Group |
| 8 | 4 | -103.53 | -94.06 | 0.262 | Spatial Coverage = Total Backtracking Duration + Adaptation to Multisensory Events + Age Group |
| 9 | 4 | -97.90 | -88.43 | 0.207 | Spatial Coverage = Total Backtracking Duration + Adaptation to Multisensory Events + Total Backtracking Duration × Adaptation to Multisensory Events |
| 10 | 4 | -109.03 | -99.55 | 0.311 | Spatial Coverage = Total Backtracking Duration + Age Group + Total Backtracking Duration × Age Group |
| 11 | 4 | -91.65 | -82.17 | 0.142 | Spatial Coverage = Adaptation to Multisensory Events + Age Group + Adaptation to Multisensory Events × Age Group |
| 12 | 5 | -103.94 | -92.10 | 0.274 | Spatial Coverage = Total Backtracking Duration + Adaptation to Multisensory Events + Age Group + Total Backtracking Duration × Adaptation to Multisensory Events |
| 13 | 5 | -107.04 | -95.20 | 0.302 | Spatial Coverage = Total Backtracking Duration + Adaptation to Multisensory Events + Age Group + Total Backtracking Duration × Age Group |
| 14 | 5 | -102.23 | -90.38 | 0.258 | Spatial Coverage = Total Backtracking Duration + Adaptation to Multisensory Events + Age Group + Adaptation to Multisensory Events × Age Group |
| 15 | 6 | -111.24 | -97.02 | 0.346 | Spatial Coverage = Total Backtracking Duration + Adaptation to Multisensory Events + Age Group + Total Backtracking Duration × Adaptation to Multisensory Events + Total Backtracking Duration × Age Group |
| 16 | 6 | -104.60 | -90.38 | 0.288 | Spatial Coverage = Total Backtracking Duration + Adaptation to Multisensory Events + Age Group + Total Backtracking Duration × Adaptation to Multisensory Events + Adaptation to Multisensory Events × Age Group |
| 17 | 6 | -105.48 | -91.26 | 0.296 | Spatial Coverage = Total Backtracking Duration + Adaptation to Multisensory Events + Age Group + Total Backtracking Duration × Age Group + Adaptation to Multisensory Events × Age Group |
| 18 | 7 | -112.73 | -96.14 | 0.365 | Spatial Coverage = Total Backtracking Duration + Adaptation to Multisensory Events + Age Group + Total Backtracking Duration × Adaptation to Multisensory Events + Total Backtracking Duration × Age Group + Adaptation to Multisensory Events × Age Group |
| **19** | **8** | **-113.79** | **-94.84** | **0.381** | **Spatial Coverage = Total Backtracking Duration + Adaptation to Multisensory Events + Age Group + Total Backtracking Duration × Adaptation to Multisensory Events + Total Backtracking Duration × Age Group + Adaptation to Multisensory Events × Age Group + Total Backtracking Duration × Adaptation to Multisensory Events × Age Group** |

*Note. All models include intercept term. Null Model (Model 1) is an intercept-only model. AIC = Akaike Information Criterion; BIC = Bayesian Information Criterion; Adj. R<sup>2</sup> = Adjusted R-squared. Lower AIC/BIC values indicate better model fit.*

**Table S2.** Model Comparison for Object Coverage Analysis.

| Model | Param No. | AIC | BIC | Adj. R<sup>2</sup> | Formula |
| --- | --- | --- | --- | --- | --- |
| 2 | 2 | -143.24 | -138.50 | 0.158 | Object Coverage = Total Backtracking Duration |
| 3 | 2 | -134.51 | -129.77 | 0.059 | Object Coverage = Adaptation to Multisensory Events |
| 4 | 2 | -160.70 | -155.96 | 0.325 | Object Coverage = Age Group |
| 5 | 3 | -146.03 | -138.93 | 0.197 | Object Coverage = Total Backtracking Duration + Adaptation to Multisensory Events |
| 6 | 3 | -166.94 | -159.83 | 0.384 | Object Coverage = Total Backtracking Duration + Age Group |
| 7 | 3 | -161.18 | -154.07 | 0.337 | Object Coverage = Adaptation to Multisensory Events + Age Group |
| 8 | 4 | -167.16 | -157.68 | 0.393 | Object Coverage = Total Backtracking Duration + Adaptation to Multisensory Events + Age Group |
| 9 | 4 | -144.39 | -134.91 | 0.190 | Object Coverage = Total Backtracking Duration + Adaptation to Multisensory Events + Total Backtracking Duration × Adaptation to Multisensory Events |
| 10 | 4 | -165.14 | -155.67 | 0.377 | Object Coverage = Total Backtracking Duration + Age Group + Total Backtracking Duration × Age Group |
| 11 | 4 | -162.94 | -153.47 | 0.360 | Object Coverage = Adaptation to Multisensory Events + Age Group + Adaptation to Multisensory Events × Age Group |
| 12 | 5 | -165.18 | -153.33 | 0.385 | Object Coverage = Total Backtracking Duration + Adaptation to Multisensory Events + Age Group + Total Backtracking Duration × Adaptation to Multisensory Events |
| 13 | 5 | -165.54 | -153.69 | 0.388 | Object Coverage = Total Backtracking Duration + Adaptation to Multisensory Events + Age Group + Total Backtracking Duration × Age Group |
| **14** | **5** | **-170.99** | **-159.14** | **0.428** | **Object Coverage = Total Backtracking Duration + Adaptation to Multisensory Events + Age Group + Adaptation to Multisensory Events × Age Group** |
| 15 | 6 | -163.54 | -149.33 | 0.379 | Object Coverage = Total Backtracking Duration + Adaptation to Multisensory Events + Age Group + Total Backtracking Duration × Adaptation to Multisensory Events + Total Backtracking Duration × Age Group |
| 16 | 6 | -169.88 | -155.66 | 0.427 | Object Coverage = Total Backtracking Duration + Adaptation to Multisensory Events + Age Group + Total Backtracking Duration × Adaptation to Multisensory Events + Adaptation to Multisensory Events × Age Group |
| 17 | 6 | -169.17 | -154.96 | 0.422 | Object Coverage = Total Backtracking Duration + Adaptation to Multisensory Events + Age Group + Total Backtracking Duration × Age Group + Adaptation to Multisensory Events × Age Group |
| 18 | 7 | -168.44 | -151.85 | 0.423 | Object Coverage = Total Backtracking Duration + Adaptation to Multisensory Events + Age Group + Total Backtracking Duration × Adaptation to Multisensory Events + Total Backtracking Duration × Age Group + Adaptation to Multisensory Events × Age Group |
| 19 | 8 | -166.58 | -147.62 | 0.416 | Object Coverage = Total Backtracking Duration + Adaptation to Multisensory Events + Age Group + Total Backtracking Duration × Adaptation to Multisensory Events + Total Backtracking Duration × Age Group + Adaptation to Multisensory Events × Age Group + Total Backtracking Duration × Adaptation to Multisensory Events × Age Group |

Note. All models include intercept term. Null Model (Model 1) is an intercept-only model. AIC = Akaike Information Criterion; BIC = Bayesian Information Criterion; Adj. R<sup>2</sup> = Adjusted R-squared. Lower AIC/BIC values indicate better model fit.

**Table S3.** Model Comparison for Target Search Success Rate Analysis.

| Model | Param No. | AIC | BIC | Adj. R<sup>2</sup> | Formula |
| --- | --- | --- | --- | --- | --- |
| 2 | 2 | -55.58 | -50.98 | 0.151 | Target Search Success Rate = Total Backtracking Duration |
| 3 | 2 | -43.43 | -38.82 | -0.000 | Target Search Success Rate = Adaptation to Multisensory Events |
| 4 | 2 | -50.81 | -46.21 | 0.095 | Target Search Success Rate = Age Group |
| 5 | 3 | -53.87 | -46.96 | 0.143 | Target Search Success Rate = Total Backtracking Duration + Adaptation to Multisensory Events |
| 6 | 3 | -58.82 | -51.91 | 0.198 | Target Search Success Rate = Total Backtracking Duration + Age Group |
| 7 | 3 | -48.92 | -42.00 | 0.083 | Target Search Success Rate = Adaptation to Multisensory Events + Age Group |
| 8 | 4 | -56.83 | -47.61 | 0.187 | Target Search Success Rate = Total Backtracking Duration + Adaptation to Multisensory Events + Age Group |
| 9 | 4 | -51.88 | -42.66 | 0.130 | Target Search Success Rate = Total Backtracking Duration + Adaptation to Multisensory Events + Total Backtracking Duration × Adaptation to Multisensory Events |
| **10** | **4** | **-61.66** | **-52.44** | **0.238** | **Target Search Success Rate = Total Backtracking Duration + Age Group + Total Backtracking Duration × Age Group** |
| 11 | 4 | -47.30 | -38.09 | 0.075 | Target Search Success Rate = Adaptation to Multisensory Events + Age Group + Adaptation to Multisensory Events × Age Group |
| 12 | 5 | -54.89 | -43.37 | 0.175 | Target Search Success Rate = Total Backtracking Duration + Adaptation to Multisensory Events + Age Group + Total Backtracking Duration × Adaptation to Multisensory Events |
| 13 | 5 | -59.70 | -48.18 | 0.227 | Target Search Success Rate = Total Backtracking Duration + Adaptation to Multisensory Events + Age Group + Total Backtracking Duration × Age Group |
| 14 | 5 | -54.95 | -43.43 | 0.176 | Target Search Success Rate = Total Backtracking Duration + Adaptation to Multisensory Events + Age Group + Adaptation to Multisensory Events × Age Group |
| 15 | 6 | -58.10 | -44.27 | 0.220 | Target Search Success Rate = Total Backtracking Duration + Adaptation to Multisensory Events + Age Group + Total Backtracking Duration × Adaptation to Multisensory Events + Total Backtracking Duration × Age Group |
| 16 | 6 | -53.12 | -39.30 | 0.166 | Target Search Success Rate = Total Backtracking Duration + Adaptation to Multisensory Events + Age Group + Total Backtracking Duration × Adaptation to Multisensory Events + Adaptation to Multisensory Events × Age Group |
| 17 | 6 | -58.07 | -44.25 | 0.220 | Target Search Success Rate = Total Backtracking Duration + Adaptation to Multisensory Events + Age Group + Total Backtracking Duration × Age Group + Adaptation to Multisensory Events × Age Group |
| 18 | 7 | -56.26 | -40.14 | 0.210 | Target Search Success Rate = Total Backtracking Duration + Adaptation to Multisensory Events + Age Group + Total Backtracking Duration × Adaptation to Multisensory Events + Total Backtracking Duration × Age Group + Adaptation to Multisensory Events × Age Group |
| 19 | 8 | -54.95 | -36.52 | 0.206 | Target Search Success Rate = Total Backtracking Duration + Adaptation to Multisensory Events + Age Group + Total Backtracking Duration × Adaptation to Multisensory Events + Total Backtracking Duration × Age Group + Adaptation to Multisensory Events × Age Group + Total Backtracking Duration × Adaptation to Multisensory Events × Age Group |

*Note. All models include intercept term. Null Model (Model 1) is an intercept-only model. AIC = Akaike Information Criterion; BIC = Bayesian Information Criterion; Adj. R<sup>2</sup> = Adjusted R-squared. Lower AIC/BIC values indicate better model fit.*

**Supplementary Analysis: Fixation on the sound source increased the mean stop duration**

The analysis in the main text only considered behavior in response to the onset of the sound. However, an individual’s stop duration was likely related to their ability to locate the sound source. Therefore, to test how the localization of the sound source affected learning and behavior, we investigated the relationship between fixation on the source of the sound and stop duration. Fixation status (source fixated or not-fixated) was added to the linear mixed model predicting average stop duration from age group and number of encounters. For simplicity, this model only included multisensory events. (A separate model of only control events found no significant differences, *p*s > .137). As illustrated in Figure 6B, the model showed a significant main effect of fixation (*β* = 2.23, *z* = 7.1, *p* < .001), with fixated events associated with a 2.23-second increase in stop duration. Critically, we found a three-way interaction of age group × fixation × encounter number (*β* = 0.55, *z* = 2.08, *p* = .025). This interaction indicated that adults had longer initial stop durations that decreased more over subsequent encounters when the source was fixated. This again suggests that adult stopping durations indicated depth of learning about an object and that learning was magnified, leading to faster adaptation on next encounters, when the source object was fixated. There was a trend towards the same effect in children, but it appears that they did not learn as much from their first encounters, reducing adaptation on subsequent encounters with the same objects.

## Footnotes

1 Degrees of freedom were estimated using the Kenward-Roger approximation to account for the variance-covariance structure in the linear mixed-effects models.

